# CDK8 is a critical effector of cell fate dysregulation in *SF3B1*-mutant MDS

**DOI:** 10.1101/2025.08.14.670174

**Authors:** Elizabeth A. Bonner, Tun-Yun Hsueh, Axia Song, Laura Baquero Galvis, Erica A. Arriaga-Gomez, Rasika Venkataraman, Sayantani Sinha, Evan J. Nguyen, Paul Brent Ferrell, Robert S. Welner, Rui Lu, H. Joachim Deeg, Derek L. Stirewalt, Sergei Doulatov, Stanley C. Lee

## Abstract

Mutations in RNA splicing factor *SF3B1* are among the most common in MDS and are strongly associated with MDS with ring sideroblasts (MDS-RS). While aberrant splicing of terminal erythroid regulators has been implicated in MDS pathogenesis, the impact of *SF3B1* mutations on early hematopoietic progenitor function remains unclear. Here, we identify CDK8, a key kinase of the mediator complex involved in transcriptional regulation, as a recurrent mis-spliced target in *SF3B1*-mutant MDS. Mutant SF3B1 induces cryptic 3′ splice site selection in CDK8, leading to loss of CDK8 mRNA and protein. Using primary human HSPCs, our study identifies CDK8 as an important regulator of HSPC homeostasis and cell fate determination. CDK8 depletion results in expansion of HSPCs and shifts differentiation toward the erythroid and myeloid lineages, mirroring phenotypes observed in *SF3B1*-mutant MDS. Lastly, functional restoration of CDK8 rescues early erythroid phenotypes in *SF3B1*-mutant cells. These findings implicate CDK8 mis-splicing as a mechanistic driver of altered progenitor fate and dysplasia in *SF3B1*-mutant MDS, linking aberrant splicing to transcriptional dysregulation and hematopoietic lineage commitment.

## INTRODUCTION

Myelodysplastic syndromes (MDS) are a heterogenous group of age-related clonal myeloid disorders characterized by cytopenia, morphologic dysplasia, and an increased risk of developing secondary acute myeloid leukemia (AML). The etiology of MDS can be broadly attributed to the complex interplay between genetic, epigenetic and environmental factors that cause functional abnormalities in bone marrow resident hematopoietic stem cells (HSCs), leading to clonal outgrowth and ineffective hematopoiesis. Somatic mutations in the RNA splicing factor *SF3B1* are highly prevalent in MDS, occurring in ∼25% of patients, and in over 70% of patients with ring sideroblasts (MDS-RS) [1–6]. SF3B1 is a member of the U2 small nuclear ribonucleoprotein (U2 snRNP) complex, which is responsible for intronic branchpoint recognition, an essential step in pre-mRNA splicing. MDS-associated *SF3B1* mutations occur in hot-spot regions within the heat domains, and confer neomorphic activity that commonly results in cryptic branchpoint selection and the preferential usage of intron-proximal 3’ splice sites [7–12]. Aberrant splicing of transcripts such as ABCB7, TMEM14C, MAP3K7, EFRE and PPOX by mutant SF3B1 have been directly linked to impaired terminal erythroid maturation, deregulation of iron homeostasis and ring sideroblast formation, which are prominent features of MDS-RS.[9, 13–16]

In addition to alterations in cellular splicing repertoires, RNA sequencing has demonstrated that *SF3B1* mutations cause alterations to the global transcriptional landscape of hematopoietic stem and progenitor cells (HSPCs). Single cell transcriptomic analysis of *SF3B1*-mutant CD34^+^ HSPCs shows that the cell transcriptional states of mutant HSPCs largely accumulate in myeloid and erythroid progenitor compartments and may be primed for preferential differentiation along the erythroid lineage.[17, 18] While many individual mis-spliced targets have been linked to deficits in late erythroid development, how mutant *SF3B1* disrupts the function of early hematopoietic progenitors remains unclear. Since a vast body of evidence has shown that lineage commitment is heavily reliant on appropriate gene expression to facilitate hematopoietic differentiation,[19–23] we hypothesize that *SF3B1* mutations may be disrupting transcriptional cofactors responsible for fine tuning gene expression.

In this study, we discovered that CDK8, a member of the mediator kinase module, is consistently mis-spliced in *SF3B1*-mutant MDS. CDK8 is the catalytic component of mediator kinase module, an accessory subunit of mediator complex that bridges transcription factors with the RNA polymerase II (RNAPII) machinery and integrates various cellular signals to modulate transcriptional output.[24] Initially thought to function as a transcriptional repressor through phosphorylation of the RNAPII C-terminal domain,[25, 26] published work indicates CDK8 functions as a transcriptional activator in the hypoxia response network and p53 pathways.[27, 28] Additionally, CDK8 kinase activity has been shown to be an important modulator of WNT, TGFβ, and INF𝛾 signaling pathways by phosphorylating downstream effectors such as c-MYC, SMADs, STAT1 and STAT3.[29–36] Given the pleiotropic roles of CDK8 as a transcriptional regulator, we investigated how MDS-associated *SF3B1* mutations promote CDK8 deregulation and its impact on dysplastic hematopoietic differentiation in MDS.

## METHODS

### Antibodies and oligos

All antibody information is available in **Supplemental Table S1**. All primers used in this study are available in **Supplemental Tables S2**.

### Cell Culture

Human umbilical cord blood CD34^+^ cells (STEMCELL Technologies) and GCSF-mobilized adult CD34^+^ cells (Fred Hutch Co-Operative Center for Excellence on Hematology (CCEH)) were thawed according to the supplier’s instructions. Cells were cultured in SFEM-II (STEMCELL Technologies) supplemented with recombinant human cytokines (50 ng/mL SCF, 50 ng/mL FLT3L, 50 ng/mL TPO; all from Peprotech), 10 μg/mL LDL (STEMCELL Technologies), 500 nM SR1 (Selleck Chem.), 1 μM UM729 (STEMCELL Technologies) for 24 hours prior to nucleofection. K562, 293T, HNT34, and Nalm6 cells were cultured in base media (IMDM, DMEM, and RPMI, respectively) supplemented with penicillin-streptomycin (100 U/mL, Gibco, No. 15140122), Glutamax (1%, Gibco, No. 35050061) and heat-inactivated FBS (10%: K562, 293T, NALM6, 20%: HNT34). The MDS-RS patient derived *SF3B1*-G742D and isogenic *SF3B1*-WT iPSC-derived hematopoietic progenitors (iPSC-HPCs) were generated and maintained as previously described.[37] Briefly, cells were maintained in SFEM-II (STEMCELL Technologies) supplemented with penicillin-streptomycin (100 U/mL, Gibco, No. 15140122), doxycycline (2 µg/mL) and cytokines (10 ng/mL IL-3, 50 ng/mL IL-6, 50 ng/mL TPO, 50 ng/mL SCF, 50 ng/mL FLT3L). Cells were washed and replated at a density of 5-7x10^5^ cells/ mL every 3 days and maintained below 1.5x10^6^ cells/mL. The iPSC-HPCs were maintained in culture for no more than 7 weeks. All cell lines were maintained in a 37°C/5% CO_2_ incubator.

### Gene Editing

Nucleofection of CD34^+^ HSPCs was carried out as previously described [38] using the P3 4D Nucleofector kit (Lonza). Briefly, 24 hours post-thaw, 2-4 x10^5^ cells were collected and rinsed in PBS and re-suspended in 20 μL nucleofection media consisting of 2.5 pmol/μL Sp.Cas9 (Aldeveron), 2.5 pmol/μL guide RNA (1.25 pmol/μl each guide, Synthego), in nucleofector solution supplemented with buffer P3. Cells were nucleofected in the 4D nucleofector (Lonza) using program DS 150 for buffer P3. Cells were recovered in 200 μL complete SFEM-II (see above) and allowed to recover for 24 hours prior to use. For applications requires greater cell numbers, multiple nucleofection reactions were performed and subsequently mixed post nucleofection. Guide RNA sequences are included in **Supplemental Table S2**.

### Lentivirus production

Lentiviral particle production was performed using low-passage 293T cells (Takara ClonTech). Cells were seeded in 10 cm tissue culture dishes at a density of 3.2x10^4^ cells/cm^2^ in 10 mL complete DMEM (see above). 2 hours prior to transfection media was removed and replaced with 8 mL complete DMEM (see above). Transfection was performed 24 hours post plating by adding transfection master mix. To make the transfection master mix 12 μg of viral plasmid, and 12 μg of packaging viral mix (3:1 ratio of psPAX2:VSVG) was added to 2.34 mL Optimem (Gibco). 30 μg of polyethylenimine (Polysciences, 23966-100) was added to the media/plasmid solution and vortexed for 30 seconds. Transfection master mix was allowed to sit at room temperature for 20 minutes prior to adding it to 293T cells, drop-wise. Complete media changed was performed 16 hours post-transfection by removing media/transfection mix and replaced with 10.5 mL of complete DMEM. Supernatants containing lentiviral particles were harvested 72 hours post transfection, snap-frozen in liquid nitrogen, and stored at -80°C. To concentrate lentiviral particles, fresh viral supernatant was first filtered of cell debris in using Millex -GP syringe filters (Millipore, SLGVM33RS), and concentrated using Amicron Ultra-15 centrifugal filter units (Millipore, UFC905024), snap frozen and stored at -80°C.

### Immunoblotting

Cells were lysed in RIPA buffer (Invitrogen, CAT#89900) supplemented with protease inhibitors (ThermoFisher, CAT#78446) and sonicated (Bioruptor UCD-200, Diagenode) for 5 minutes in 30 second on/off cycles on medium setting. Protein concentrations were normalized using a Pierce BCA protein assay kit (Thermo Fisher Scientific CAT#23227). 20 ng of protean lysate was mixed with 6XSDS loading buffer (0.3M Tris-HCL, 10% SDS, 60% Glycerol, 6% 2-mercaptoethanol, 1% bromophenol blue), loaded in pre-cast SDS acrylamide gels (BioRad, CAT#4561084) and run at 90V. Blots were transferred using the Bio-Rad turbo transfer apparatus following manufacturers recommendation. Membranes were blocked in Tris-buffered saline with 0.05% Tween-20 (TBS-T) and 5% milk for 1h at room temperature and immunoblotted with primary antibodies overnight at 4°C. Immunoblot antibodies are listed in **Supplemental Table S1**. Membranes were washed 3 times with TBS-T and incubated for 1 h at room temperature with secondary antibodies conjugated to horseradish peroxidase. Blots were imaged using ECL Substrate (BioRad, CAT#1705062) using AFP mini-medical 90 film processor.

### RT-PCR

200 ng Trizol extracted RNA was used to make cDNA using iScript (BioRad, CAT# 1708891). PCR was performed using GoTAQ (Promega, CAT#M7123). PCR products were run using the Protean II xi electrophoresis tank (BioRad) in a 12% native polyacrylamide gel (BioRad, CAT#1610156) in 1XTBE (Fisher Chemical, SO-0016) at 45V for 48 hours at 4°C and imaged using ethidium bromide (Fisher Scientific, CAT#BP1302-10) on a Gel Doc XR (Bio-Rad). All RT-PCR primers are listed in **Supplemental Table S2.**

### Flow cytometry

Cells were harvested and washed in cold FACS buffer (1XPBS, 2%FBS). Cells were resuspended in 100 μL staining buffer (FACs buffer with a 1:200 dilution of each antibody) and incubated in the dark at 4C for 20 minutes. Staining buffer was washed by adding 1 mL FACS buffer and resuspended in 100-300 μL FACS buffer with 1:500 dilution dapi (Millipore, 10236276001). Flow cytometry was performed on the BD FACSymphony A5. Flow cytometry antibodies are listed in **Supplemental Table S1**.

### Primary human MDS patient samples

Cryopreserved de-identified bone marrow specimens from MDS patients were obtained from the Fred Hutchinson Cancer Center/University of Washington Hematopoietic Diseases Repository (FHCC/UW-HDR). As part of the FHCC/UW-HDR, patient specimens are collected and stored under the oversight of Fred Hutch Institution Review Office, and all participants provided written informed consent that adhered to the guidelines of 1975 Declaration of Helsinki. MDS diagnosis for all the patients was confirmed using established guidelines at the time of diagnosis. See **Supplemental Table S3** for detailed genetic information for each patient sample.

### Animals models

All animal experiments were performed with approval by and in accordance with Fred Hutchinson Cancer Center (FHCC) Institutional Animal Care and Use Committees guidelines. Animal experiments were performed within the FHCC Comparative Medicine facility. For xenotransplantation experiments, 6–7-week-old female NOD.Cg-*Prdkc*^SCID^ *Il2rψ*^-/-^ (NSG) mice (Fred Hutchinson Cancer Center Comparative Medicine Shared Resources) were used. NSG mice were sub-lethally irradiated (2.0 cGy) 24 hours prior to transplantation. Twenty-four hours post recovery from nucleofection, ∼4-8 x10^4^ cells were transplanted intravenously via tail vein injection. Peripheral blood was monitored by flow cytometry every 4 weeks to track human cell chimerism. At 20 weeks post-xenotransplantation, hematopoietic organs (bone marrow, spleen and peripheral blood) were harvested and analyzed by flow cytometry.

### *Ex vivo* differentiation assays

For colony-forming unit (CFU) assays, umbilical cord blood-derived CD34^+^ HSPCs were plated 24 hours post nucleofection in 1 mL MethoCult H4034 (STEMCELL Technologies). Cells were plated at a density of 500 cells/mL in triplicate plates. Colonies were counted and scored 11 days post-plating. For myeloid differentiation assays, healthy adult CD34^+^ HSPCs were electroporated and plated at a density of 2.5 x10^4^ cells/well in 96-well U-bottom plates in SFEM-II containing Myeloid II Expansion supplement (STEMCELL Technologies). Differentiation was monitored by flow cytometry for expression of CD14 and CD13. Erythroid differentiation was performed using healthy adult CD34+ HSPCs following previously described protocols.[39] Briefly, 1x10^5^ cells were plated in 1 mL of expansion media (SFEM supplemented with 100 ng/mL FLT3 ligand, 100ng/mL SCF, 5 ng/mL IL-3, 20 ng/mL IL-6) for 3 days. On day 4 cells were plated at a density of 1 x10^5^ cells/ mL in Stage 1 media (IMDM, 15% FBS, 2 mM glutamine, 1% BSA, 500 μg/mL holo-transferrin, 10 μg/mL insulin, 100 ng/mL SCF, 5 ng/mL IL3, 6 U/mL EPO) and cultured for 5 days. On day 9 cells were plated at a density of 2x10^5^ cells/ mL in Stage 2 media (IMDM, 15% FBS, 2 mM glutamine, 1% BSA, 500 μg/mL holo-transferrin, 10 μg/mL insulin, 100 ng/mL SCF, 6 U/mL EPO). On day 14 cells were seeded at 3 x10^5^ cells/mL in Stage 3 media (IMDM, 15% FBS, 2 mM glutamine, 1% BSA, 500 μg/mL holo-transferrin, 10 μg/mL insulin, 2 U/mL EPO). The appropriate media was supplemented in the middle of each stage to maintain cells at a density below 2 x10^6^ cells/mL. Differentiation was monitored by flow cytometry for expression of CD71 and CD235a. For erythro-myeloid differentiation of iPSC-HPCs, cells were plated at a density of 2x10^5^ cells/mL in 1 mL differentiation media (IMDM, 2mM glutamine, 15 ng/mL G-CSF, 100 ng/mL SCF, 40 ng/mL FLT3, 0.5 U/mL EPO, 10 ng/mL IL-3). Cells were maintained at a density below 5x10^5^ for 14 days. Media formulation was changed on 6 days in culture (increase [EPO] to 3 U/mL]. Differentiation was monitored by flow cytometry. All human cytokines were purchased from PeproTech.

### RNA sequencing

For cultured cell lines total RNA was isolated from 5 x10^6^ K562 SF3B1^WT^, K562 SF3B1^K700E^ or HNT34 cells using TRIzol according to the manufacturer’s instruction. For CDK8 knock-down UCB derived CD34^+^ cells, 0.2-3 x10^6^ CD34^+^ GFP^+^ cells were sorted using the Symphony S6 sorter (BD Biosciences), followed by RNA isolation by Trizol. RNA quality was determined using Agilent RNA 6000 Nano chip on the Agilent 4200 TapeStation and quantification was performed using NanoDrop spectrophotometer (Thermo Fisher Scientific). Total RNA samples with RNA integrity number (RIN) >8 was selected for Illumina sequencing library preparation. One microgram of DNA-free total RNA was used to prepare the library using Ultra II Directional RNA Library Prep Kit (New England Biolabs, No. E7760L) following manufacturer’s instructions. Paired end sequencing (150 bp) of the libraries were performed using an Illumina NovaSeq SP platform.

### Amplicon sequencing

Human CD45^+^ cells were isolated from murine bone marrow 20 weeks post-transplant using the using the BD Symphony S6 sorter (BD Biosciences). DNA was extracted using the Qiagen DNeasy blood and tissue kit (CAT#69504). PCR was performed using Platinum II hot start PCR master mix (Invitrogen, CAT#14000013). PCR products were precipitated overnight using ethanol precipitation (https://projects.iq.harvard.edu/files/hlalab/files/ethanol-precipitation-of-rna_hla.pdf) and submitted for sequencing at Azetna following submission guidelines.

### Bioinformatic Analysis

For RNA sequencing, raw .fastq files were trimmed and quality was assessed using TrimGalore (https://github.com/FelixKrueger/TrimGalore). Trimmed fastq files were aligned (GRCh38, Gencode Release 48) with STAR[40] using –quantMode GeneCounts. For publicly available MDS patient data batch correction was performed using limma[41] and differential expression for all datasets was performed using DESeq2[42] comparing wild type to mutant samples. Significant changes were determined using a cutoff of |Log2 fold change| ≥ 0.5 and p-adjusted value ≤ 0.05. For splicing analysis reads were aligned (GRCh38, Gencode Release 48) using STAR along with soft-clipping and splicing changes were quantified using rMATS-turbo 4.1.1[43] with the novel splice site detection feature enabled. To generate heat map plots of publicly available data, .JC files were imported using maser (https://github.com/DiogoVeiga/maser) and significant splicing events were determined (minimum read count =10, |PSI|≥ 0.05, FDR <0.05). A list of 162 transcriptional cofactors was curated from gene ontology (https://geneontology.org/) and filtered for unique entries. Significant splicing events were filtered using the unique transcriptional cofactor list. Z scores were calculated for splicing events within each data set to control for batch effects and unsupervised clustering was performed across all samples within each splice type. For amplicon sequencing analysis, quality and trimming was performed on raw fastq files (see above). Reads were aligned to CDK8 reference sequence (NCBI NC_000013.11) and analyzed using CRISPResso[44] using a window of 50 nucleotides surrounding the guide.

### Data Availability

RNA-sequencing data generated as part of this study were deposited in the Gene Expression Omnibus Accession Number GSE314578. RNA sequencing data from published studies were downloaded from the European Nucleotide Assembly (ENA).

## RESULTS

### Focused RNA-seq analysis of transcriptional cofactors identifies recurrent mis-splicing of CDK8 in *SF3B1*-mutant MDS patients

To identify candidate transcriptional co-factors which undergo mutant SF3B1-mediated aberrant splicing we performed splicing analysis[45] using RNA sequencing data from isogenic leukemia lines (K562 *SF3B1*-WT, K562 *SF3B1*-K700E, and HNT34, a leukemia cell line that harbors an endogenous *SF3B1*-K700E mutation), as well as publicly available MDS patient samples with and without *SF3B1* mutations (**Supplemental Table S4**) [46, 47]. We curated a list of 162 transcriptional co-factors using gene ontology[48, 49] (**Supplemental Table S5**) and looked for significant splicing events from our cohort that followed a set of criteria: 1) The splicing event must be enriched in *SF3B1* mutant cells compared to wild-type (WT), 2) the splicing event deregulates transcript stability, 3) mis-splicing must occur across different SF3B1 mutations, and 4) the splicing event is exclusive to *SF3B1* mutations. Using a delta percent spliced-in (dPSI) cut off of 0.08 and FDR <0.05, our analysis showed that, of the transcripts predicted to undergo alternative 3ʹ splice site usage, members of the mediator complex are highly enriched, including MED13L, MED6 and CDK8 (**Figure 1A**, **Supplemental Table S6**). Because alternative splice site usage can either lead to stable alternative isoform usage or can introduce a premature termination codon, we next asked if any of the transcriptional co-factors also displayed changes in transcript level by RNA sequencing (**Figure 1B**). We performed differential gene expression analysis using DESeq2[50] and found that, among the 20 predicted mis-splicing events, only three transcripts (CDK8, TAF6L, TAF2) showed significant changes in expression (adjusted p-value <0.05). Notably, CDK8 was the only mediator complex member with significantly altered transcript levels. To determine if changes in CDK8 mRNA levels were specific to the *SF3B1*-K700E mutation, we performed differential expression analysis on cells from a larger cohort of MDS patients with and without *SF3B1* mutations[46, 51–53], healthy bone marrow and *SF3B1*-WT (K562) and *SF3B1*-K700E (K562, HNT34) cell lines (**Figure 1C**). While TAF6L displayed a greater decrease in mRNA expression in *SF3B1*-mutant samples, alternative 3′ splice site usage in both patient samples and cell lines was significantly enriched in the *SF3B1*-WT cells (**Figure 1A, Supplemental Figure S1A),** which did not meet our criteria. Similarly, the mis-splicing of MED13L, TAF2, and MED6 in *SF3B1*-mutant cells did not meet our criteria, either due to a lack of significant transcript downregulation or because the resulting splice variants generated stable mRNA isoforms (**Supplemental Figure S1B-H)**.

**Figure 1.**
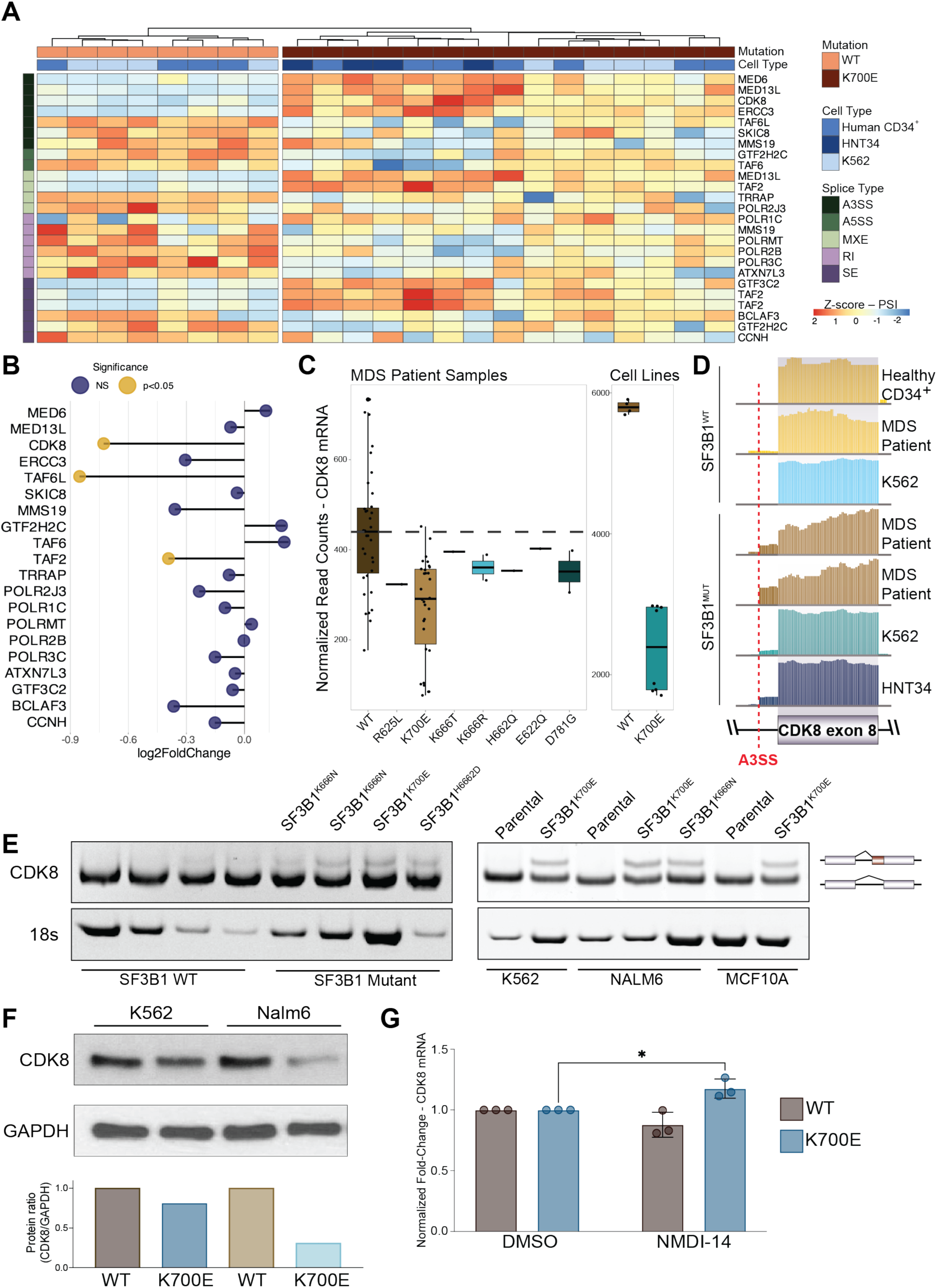
RNA sequencing reveals CDK8 deregulation in Sf3B1 mutant MDS patient samples and isogenic cell lines. **(A)** Unsupervised clustering (columns) of significant (compared to WT MDS and healthy controls) mis-spliced transcriptional cofactors (read count > 10, |PSI| > 0.08, FDR < 0.05, rows) from publicly available *SF3B1*-mutant MDS patient RNA sequencing data and isogenic cell lines (K562, HNT34). **(B)** Lollipop plot of the Log_2_ fold change (DESeq2) of significantly mis-spliced transcriptional co-factors. Yellow, adjusted pvalue < 0.05, blue, adjusted pvalue ≥ 0.05. **(C)** Normalized read counts (DESeq2) of CDK8 mRNA from publicly available data. Dashed line = mean read count of *SF3B1* WT samples. **(D)** IGV plots of the mis-spliced region of CDK8 in *SF3B1* WT and *SF3B1*-mutant MDS patient samples and cell lines. Y-axis, absolute read count. **(E)** RT-PCR and native PAGE of 158 bp surrounding the alternative 3′ splice site usage in CDK8 mRNA collected from *SF3B1*-wildtype and *SF3B1*-mutant MDS patient samples (left) and *SF3B1*-isogenic cell lines (right). 18s rRNA was used as loading control (bottom) **(F)** Western blot of CDK8 protein levels from wild type and *SF3B1* mutant cell lines **(G)** qPCR mean ± SD of isogenic K562 cell lines treated with NMDI-14 (n=3 independent experiments). Treated sample were normalized to untreated controls. *, p < 0.05, two-way ANOVA.

Due to alternative 3′ splice site usage, CDK8 is predicted to include 14-base pairs of intronic sequence proximal to exon 8. To confirm this, we examined CDK8 read coverage in the Integrated Genome Viewer[54] (IGV) and observed intronic inclusion exclusively in samples from *SF3B1*-mutant cells (**Figure 1D**). Additionally, we did not observe CDK8 mis-splicing in other common spliceosome mutations (**Supplemental Figure S1I)** Taken together, these data indicate that *SF3B1* mutations affect only a small subset of transcriptional co-factors, and among those, CDK8 is the only one consistently showing decreased mRNA transcript levels across different *SF3B1*-mutant contexts. (**Supplemental Figure S1A-D**)

### Mutant SF3B1-induced mis-splicing of CDK8 results in loss of expression via nonsense-mediated decay

We next sought to validate CDK8 mis-splicing in *SF3B1*-mutant MDS patient samples and cell lines. We designed PCR primers which spanned exon-exon boundaries around CDK8 exon 8 and performed RT-PCR followed by native polyacrylamide gel electrophoresis (PAGE). We detect the 14 base-pair intronic sequence inclusion in *SF3B1*-mutant but not *SF3B1*-WT MDS patient samples (**Supplemental Table S3**) and isogenic cell lines (**Figure 1E**). We then determined if reductions in CDK8 mRNA corresponded to a reduction of CDK8 at the protein level. We performed western blot in *SF3B1*-mutant cell lines and found that CDK8 protein was reduced in *SF3B1* mutant samples but not their WT counterparts (**Figure 1F**). The inclusion of 14 base pairs of intronic sequence is predicted to lead to a frameshift, resulting in reduced CDK8 mRNA through the quality surveillance pathway nonsense-mediated mRNA decay (NMD). To verify this prediction, we treated cells with an NMD inhibitor, NMDi-14, and performed qPCR in isogenic *SF3B1*-mutant cell lines. We found that NMDi-14 treatment restored CDK8 mRNA levels in *SF3B1*-mutant cells but not in *SF3B1*-WT cells (**Figure 1G**). Together, these data verify that CDK8 is mis-spliced by mutant SF3B1 leading to reduction of CDK8 mRNA and protein due to mRNA degradation by NMD.

### Loss of CDK8 causes defective hematopoietic stem and progenitor cell function and multi-lineage differentiation *in vitro* and *in vivo*

Phenotypically, *SF3B1*-mutant MDS patients present with an erythroblasts expansion combined with ineffective erythropoiesis, and single cell transcriptomic data indicate that *SF3B1*-mutant HSPCs have an accumulation of cell states resembling early erythroid and myeloid precursors.[13, 17, 18, 55, 56] To determine if CDK8 loss is contributing to the shift in frequency of early myeloid and erythroid progenitors, we used a dual-guide RNA-based CRISPR/Cas9 system[57] to target CDK8 (CDK8^KO^) or the safe harbor locus AAVS1 (AAVS1^KO^) locus as negative control in human CD34^+^ HSPCs (**Supplemental Figure S2A-C**). To monitor differentiation along a single lineage, we used G-CSF-mobilized CD34^+^ cells from healthy adults and performed liquid culture differentiation along the myeloid or erythroid axis. We observed that CDK8^KO^ HSPCs produced a higher percentage of CD13^+^CD14^+^ myeloid cells, and CD71^+^CD235a^+^ erythroid cells compared to AAVS1^KO^ cells (**Figure 2A-B**). To monitor multi-lineage differentiation of early progenitors, we generated CDK8^KO^ cells from umbilical cord blood (UBC)-derived CD34^+^ HSPCs and performed colony-forming unit (CFU) assays (**Figure 2C**). Our data showed that CDK8^KO^ HSPCs generated significantly more colonies, driven by a marked increase in CFU-Granulocyte/Macrophage (CFU-G/M) colonies. While CDK8^KO^ HSPCs did not produce a significant increase in the number of early burst-forming erythroid colonies (BFU-E), total colonies collected at the end of the experiment showed that colonies derived from CDK8^KO^ HSPCs appeared more hemoglobinzed relative to control (**Figure 2D**). Together, these results indicate that CDK8 loss leads to increased differentiation along the myeloid and erythroid axes and supports our hypothesis that CDK8 mis-splicing may be contributing to the expansion of early erythroid and myeloid progenitors in *SF3B1*-mutant MDS.

**Figure 2.**
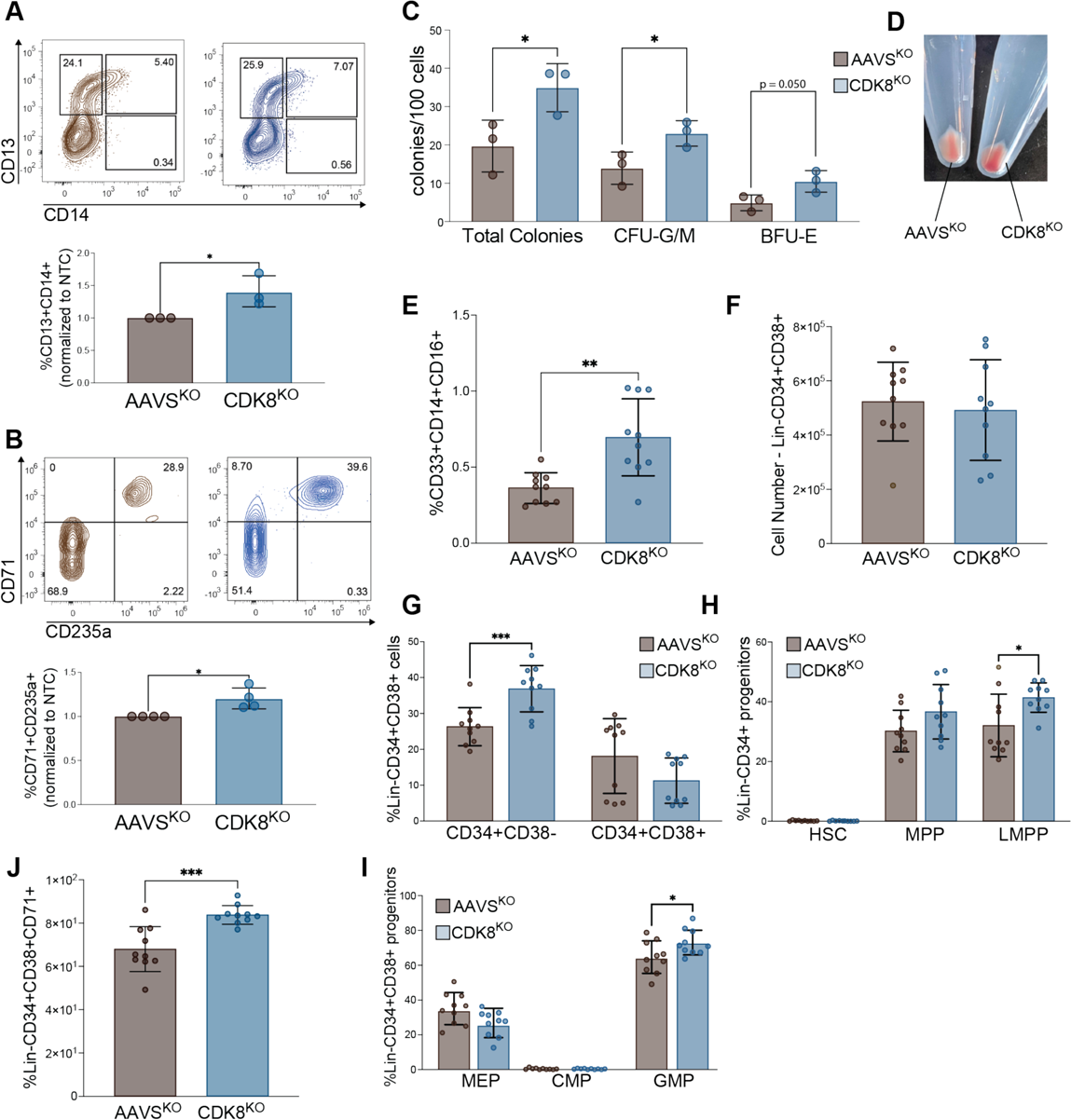
Loss of CDK8 alters human HSPC differentiation. **(A)** Representative flow plots (top) and mean ± SD (bottom) of CD13^+^CD14^+^ myelo-monocytic cells produced by GCSF-mobilized CD34^+^ cells electroporated with guide RNAs targeting AAVS1 or CDK8. **(B)** Representative flow plots (top) and mean ± SD (bottom) of CD71^+^CD235a^+^ erythroid cells produced by GCSF-mobilized CD34^+^ cells electroporated with AAVS1 or CDK8 guide RNAs. Data in (A, B) are normalized to AAVS1^KO^ group. **(C)** Number of colony forming units (CFU) and the types of colonies produced (G/M, granulocyte/macrophage progenitors, BFU-E, burst-forming unit – erythroid). Data are represented as CFU number per 100 CD34^+^ UCB cells electroporated with AAVS1 or CDK8 guide RNAs. **(D)** Representative image of CFU colony pellet following collection at assay termination. *, p < 0.05, unpaired two-tailed Student’s t-test. **(E)** Percentage of human CD45^+^CD33^+^CD14^+^CD16^+^ intermediate monocytes in the bone marrow of mice (mean ± SD). **(F)** Viable cell number of lineage negative CD34^+^CD38^+^ cells in the bone marrow. **(G)** Percentage lineage-negative (Lin^-^) CD34^+^CD38^-^ cells in bone marrow (mean ± SD). **(H)** Percentage Lin^-^CD34^+^CD38^-^ of hematopoietic stem cells (HSC, CD90^+^CD45RA^-^), multipotent progenitors (MPP, CD90^-^CD45RA^-^), and lymphoid myeloid primmed progenitors (LMPP, CD90^-^CD45RA^+^) cells in bone marrow (mean ± SD). **(I)** Percentage Lin^-^CD34^+^CD38^+^ megakaryocyte erythroid progenitors (MEP, CD45RA^-^CD123^-^), common myeloid progenitors (CMP, CD45RA^-^CD123^+^), and granulocyte macrophage progenitors (GMP, CD45RA^+^CD123^+^) in the bone marrow. **(J)** Viable cell number of Lin^-^CD34^+^CD38^+^CD71^+^ cells in the bone marrow.

To determine the long-term *in vivo* consequences of CDK8 loss in hematopoiesis, we used the same CRISPR/Cas9 system to knockout CDK8 in UCB CD34^+^ cells followed by xenotransplantation in NOD/SCID/ILR2𝛾-null (NSG) mice (**Supplemental Figure S3A**). At terminus (20 weeks post xenotransplantation), we harvested spleen, bone marrow and peripheral blood to assess lineage composition and confirm knock-down in bone marrow CD45^+^ cells (**Supplemental Figure S2C-D**). CDK8^KO^ UCB mice had a mild increase in CD33^+^ myeloid cells in the blood and bone marrow (**Supplemental Figure S3C-E**). There were no significant differences between the level of human cell chimerism (%CD45^+^) in the bone marrow mononuclear cell fraction of mice transplanted with AAVS1^KO^ or CDK8^KO^ HSPCs (**Supplemental Figure S3F**). However, there was a significant increase in CD14^+^CD16^+^ intermediate monocytes in the bone marrow (**Figure 2E**). Further examination of the bone marrow HSPC fraction (defined by CD45^+^lineage^-^CD34^+^) revealed that there was no significant increase in the total number of HSPCs between the AAVS1^KO^ and CDK8^KO^ groups (**Figure 2F**). Further, there was a mild but significant increase in the absolute frequency of the more immature progenitors (defined by CD45^+^lineage^-^CD34^+^CD38^-^) and lymphoid-primed multipotent progenitors (LMPPs; defined by CD45^+^lineage^-^CD34^+^CD38^-^CD45RA^+^CD90^-^) (**Figure 2G-H, Supplemental Figure S3G**). When we looked within the more differentiated myeloid progenitor fraction (defined by CD45^+^lineage^-^CD34^+^CD38^+^), we observed a significant increase in the relative frequency of granulocyte-macrophage progenitors (GMPs; CD45^+^lineage^-^CD34^+^CD38^+^CD45RA^+^CD123^+^) (**Figure 2I)**. While we did not see an expansion in immunophenotypic megakaryocyte-erythroid progenitors (MEPs; CD45^+^lineage^-^CD34^+^CD38^+^CD45RA^-^CD123^-^), we observed a significant expansion in a distinct subset of cells within the CD34^+^CD38^+^ compartment that express CD71 (**Figure 2J**), a population with the potential to differentiate into both erythroid and granulocytic lineages.[58, 59] Our *in vivo* data indicates that loss of CDK8 leads to an expansion of specific primitive HSPC compartments that skew towards the erythroid and granulocytic/myeloid differentiation trajectories.

### CDK8 loss deregulates hemostatic- and lineage-specific transcriptional pathways in HSPCs

Because CDK8 functions as transcriptional modulator, we next sought to determine the changes resulting from CDK8 loss in UBC CD34^+^ HSPCs. We performed RNA sequencing in CD34^+^ UCB cells treated with 2 independent short-hairpin RNAs (shRNAs) targeting CDK8 (shCDK8) or a non-targeting control (shNTC) (**Figure 3A**, **Supplemental Figure S4A-C**). We performed differential expression analysis (DESeq2) and filtered for significant (adjusted p-value <0.05) changes shared across both hairpins (**Figure 3B**, solid dots). CDK8-deficient CD34^+^ cells had increased expression of known markers of myeloid differentiation (CD14, CECAM6, S100A8, S100A9), and decreased expression in signaling intermediates of key intracellular signaling pathways including the MAP kinase signaling (FOS, FOSB, JUN, DUSP2), JAK/STAT pathway (SOCS3) and GPCR signaling (RGS1, RGS2) pathways (**Figure 3A**, **Supplemental Figure S4D**, **Supplemental Table S7**). We then asked if we could detect similar patterns of deregulation in the MDS patient cohort. We found that most of the transcripts deregulated in CDK8-deficient CD34^+^ HSPCs showed similar expression patterns in *SF3B1*-mutant cells from MDS patients (**Figure 3C**). Taken together, these results indicate that CDK8 loss leads to deregulation of several important transcriptional networks, likely resulting in the upregulation of genes associated with lineage differentiation.

**Figure 3.**
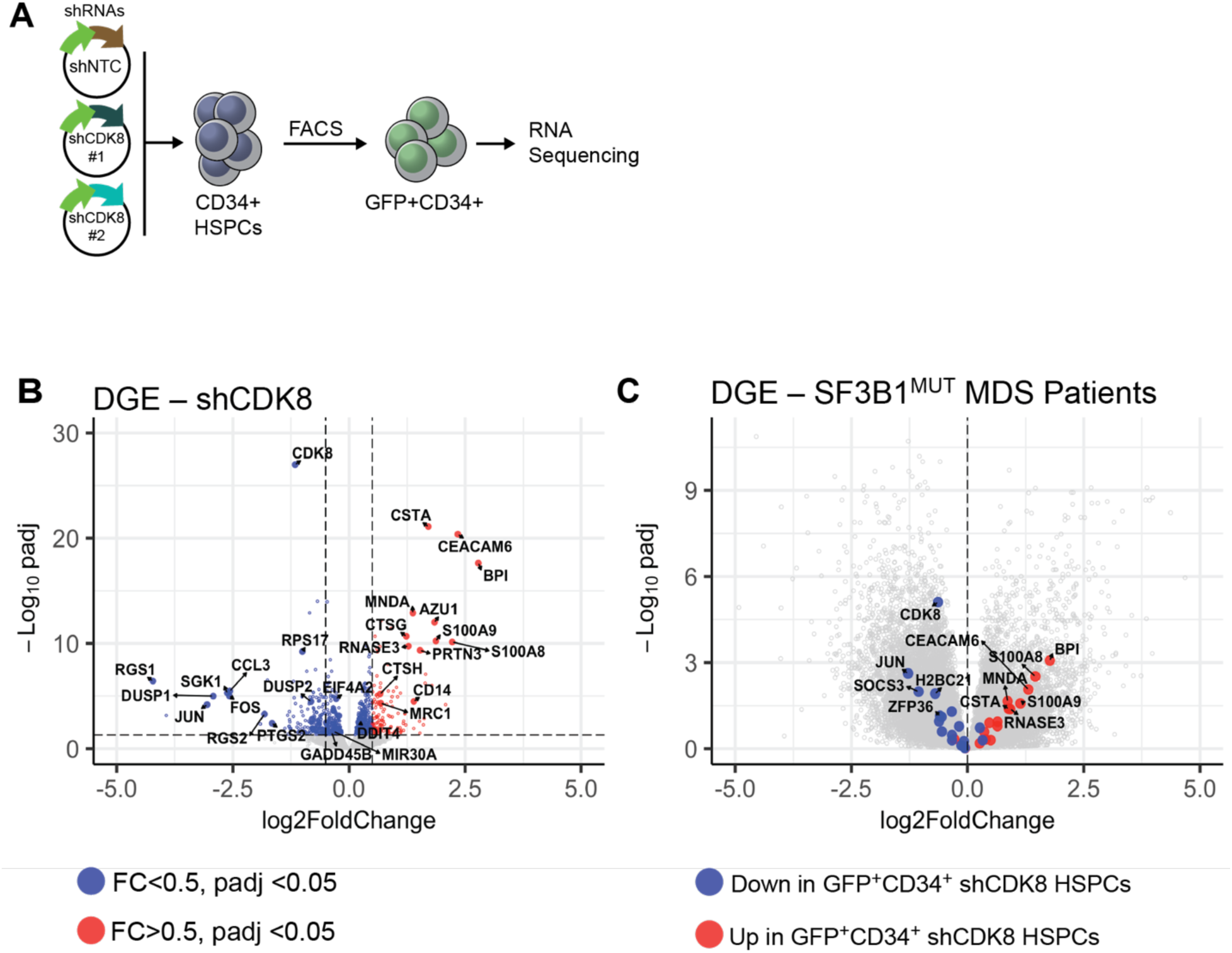
Homeostatic and lineage pathways are deregulated by CDK8 loss. **(A)** Experimental overview for RNA sequencing experiment. **(B)** Volcano plot representation of differentially expressed genes (DGE) from CDK8 knock-down cells compared to non-targeting control, showing the magnitude of difference (log_2_ fold change, x-axis) and significance (-log_10_(adjusted p-value), y-axis). Red dots, adjusted p-value < 0.05, LFC >0.5, blue dots, adjusted p-value < 0.05, LFC <0.5, solid dots, significant genes across both shRNAs. Dashed lines mark the cutoff for significance (horizontal, adjusted p-value < 0.05) and fold change (vertical, -0.5 < LFC >0.5). **(C)** Volcano plot representation of differentially expressed genes (DGE) from *SF3B1*-mutant patient samples compared to *SF3B1*-WT MDS and healthy controls. Dots, genes shown to be significantly up (red) or down (blue) in CDK8 knockdown CD34^+^ HSPCs, grey circles, all other genes.

### Restoration of CDK8 partially rescues erythroid differentiation defects in *SF3B1*-mutant iPSC model

Next, we sought to formally demonstrate the causal link between the phenotypic consequences of CDK8 loss to *SF3B1* mutations in MDS. To address this, we used a previously established isogenic pair of induced pluripotent stem cell-derived hematopoietic progenitor cell lines (iPSC-HPCs) generated from an MDS-RS patient with an *SF3B1*-G742D mutation[60] (**Supplemental Figure S5A**). We also observed in the iPSC-HPC model that CDK8 is robustly mis-spliced and downregulated at both the mRNA and protein level in *SF3B1*-mutant cells relative to their *SF3B1*-WT isogenic counterparts (**Figure 4B-C, Supplemental Figure S5B**). Consistent with prior erythroid differentiation data, we observed that *SF3B1*-mutant iPSC-HPCs have increased proportion of CD71^+^CD235a^+^ cells (**Supplemental Figure S5C)**. We therefore hypothesized that this early expansion of erythroid progenitors may be due to loss of CDK8. To determine if restoration of CDK8 could rescue erythroid differentiation defects in *SF3B1*-mutant cells, we stably introduced CDK8 cDNA or an empty vector (EV) control in *SF3B1*-mutant iPSC-HPCs and performed erythroid differentiation (**Figure 4A & 4D**). Compared to EV-transduced *SF3B1*-mutant cells, re-expression of CDK8 significantly reduced the expansion of CD71^+^CD235a^+^ cells in both erythroid differentiation conditions (**Figure 4E & 4F**) and erythro-myeloid differentiation conitions **(Supplemental Figure S5E-F)**. Taken together, our data indicates that the aberrant erythroid expansion and differentiation phenotype characteristic of *SF3B1* mutations in MDS-RS is at least partly mediated through mis-splicing and the subsequent loss of CDK8 function.

**Figure 4.**
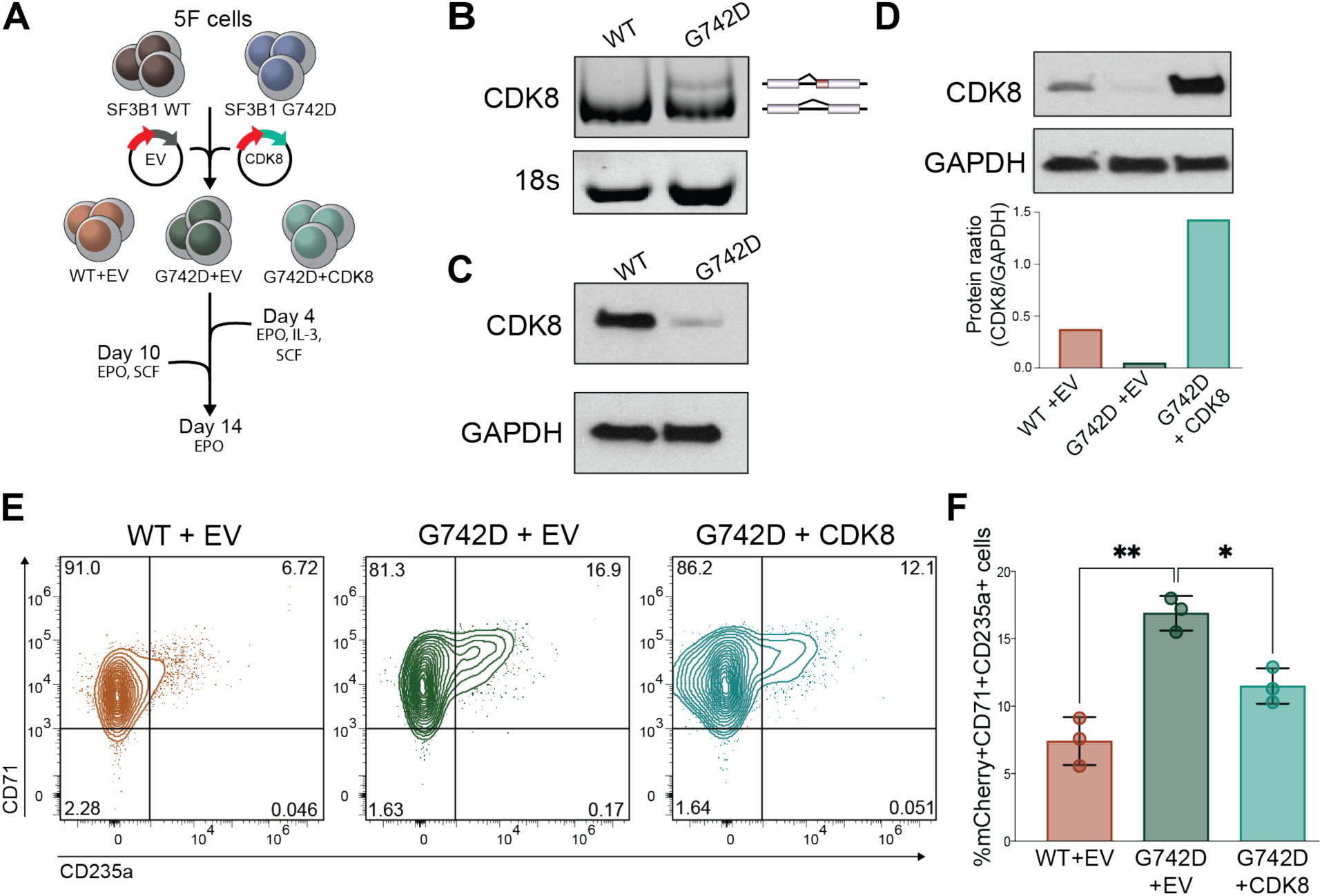
CDK8 re-expression rescues early erythroid differentiation. **(A)** Experimental overview for erythroid differentiation assay using the iPSC-HPC model. **(B)** RT-PCR and native PAGE of 158 bp surrounding the alternative 3′ splice site usage in CDK8 mRNA collected from isogenic *SF3B1* wildtype (WT) and *SF3B1-*G742D iPSC-HPCs. 18s rRNA was used as a loading control (bottom) **(C)** Western blot of CDK8 protein levels from *SF3B1* wildtype and *SF3B1* mutant iPSC-HPCs. **(D)** Western blot (top) and quantification of iPSC-HPCs transduced with lentiviral particles containing empty vector (EV), or full-length CDK8 cDNA. Western blot was quantified and protein ratio (relative to loading control, ImageJ) was plotted (bottom). **(E)** Representative flow plots and **(F)** mean ± SD (bottom) of CD71^+^CD235a^+^ iPSC-HPCs transduced with CDK8 cDNA or EV control on day 4 of erythroid differentiation. *, p < 0.05, **, p < 0.01, one-way ANOVA.

## DISCUSSION

*SF3B1* mutations are purported as functional drivers of expansion in clonal hematopoiesis and MDS. However, despite extensive research, the mechanistic understanding of how splicing defects confer a competitive advantage and skew early HSPC cell fate choice remains elusive. Here, we utilized publicly available patient transcriptomics data,[46, 51–53] *ex vivo* and *in vivo* models to identify CDK8 mis-splicing as a potential mechanism governing impaired hematopoiesis. Our findings highlight CDK8 as a critical effector in *SF3B1*-mutated MDS and elucidate its importance in regulating proper hematopoietic differentiation.

*SF3B1* is one of the most commonly mutated genes and the most frequently mutated spliceosome component in MDS and is associated with widespread splicing deregulation. [1–6, 8–12, 51, 61–64] Here, we identify CDK8 as a recurrent mis-splicing target in *SF3B1*-mutant MDS patients across a range of hotspot mutations. Mis-splicing led to deregulation of CDK8 transcript levels in mutant cells due to the introduction of a premature termination codon, resulting in CDK8 degradation by the quality surveillance pathway nonsense-mediated mRNA decay.

Previous studies have linked *SF3B1* mutations to impaired terminal erythroid differentiation, iron metabolism and the formation of ring sideroblasts,[13, 16, 51, 62] but a mechanistic link between clonal advantage and early fate choice of HSPCs had not been made. Here, we show that, as a result of CDK8 loss, primitive HSPCs compartments expand *in vivo* and may preferentially differentiate along the myeloid and erythroid axes *in vitro* and *in vivo*. This skewed differentiation parallels the expansion of immature erythroid and myeloid precursors observed in *SF3B1*-mutant MDS patients and from *SF3B1*-mutant HSPCs. Thus, our findings provide a mechanistic link between *SF3B1* mutations and disrupted early HSPC fate via partial loss of CDK8.

CDK8 is a context-specific transcriptional regulator in solid tumors, influencing pathways like WNT, p53 and IFN-STAT signaling.[29–36] Using RNA sequencing, our study identifies CDK8 as an important regulator of myeloid marker gene expression and indicates that deregulation of MYC and STAT pathways may be driving altered HSPC cell fate choice in *SF3B1*-mutant cells. Further supporting our findings, published work has shown that both MYC and JAK/STAT pathways play important regulatory roles in HSPC self-renewal, lineage commitment, and leukemogenic transformation.[65–71]

*SF3B1*-mutant MDS is generally considered low-risk, with current treatments focused on managing anemia. Our data shows that reintroduction of CDK8 rescues erythroid abnormalities in *SF3B1*-mutant cell models and suggests a targeted approach to correcting differentiation defects, potentially through the use of splice correcting antisense oligonucleotides (ASOs). Additionally, the observation that CDK8 loss enhances erythroid output underscores its therapeutic relevance, particularly given the abundance of available CDK8 kinase inhibitors that could be repurposed to promote erythropoiesis in other anemic conditions.

## ACKNOWLEDGEMENTS

The authors would like to thank the S. Lee lab for discussion; Daniel Kuppers (FHCC), Anthony Rongvaux (FHCC) and Stefan Radtke (FHCC) for technical advise; Sean Bennette (FHCC) for advice on bioinformatics analysis; Tim Monahan, Bella Morocho and the Fred Hutch Cancer Center/University of Washington Hematopoietic Diseases Repository (FHCC/UW-HDR) for MDS samples. S.C. Lee is supported by grants from the National Cancer Institute (R01 CA292932), National Institute of Diabetes and Digestive and Kidney Diseases (RC2 DK127989), an Edward P. Evans Foundation Discovery Research Grant, a Scholar Award and a Bridge Grant from the American Society of Hematology, and a Mark Foundation for Cancer Research Endeavor Award. R. Venkataraman is supported by an American Society of Hematology Graduate Hematology Award; S. Sinha is supported by an American Cancer Society Postdoctoral Fellowship (PF-23-1145387-01-ET). PBF is supported by a Mark Foundation for Cancer Research Endeavor Award, a 350 Novartis Global Scholar Award, the National Institute of Diabetes and Digestive and Kidney Diseases (R56 DK138826), VA MERIT Award 351 (I01BX005991). R.S. Welner is support by the National Heart, Lung, and Blood Institute (P01 HL131477), a Mark Foundation for Cancer Research Endeavor Award, and an Edward P. Evans Foundation Discvoery Research Grant. R. Lu is supported by a Mark Foundation for Cancer Research Endeavor Award, the grants from the National Cancer Institute (R01 CA259480), American Cancer Society (RSG-22-036-01-DMC), and the Gabrielle’s Angel Foundation for Cancer Research. S.D. is supported by the grants from National Heart, Lung, and Blood Institute (R01 HL151651 and R01 HL169156), National Institute of Diabetes and Digestive and Kidney Diseases (RC2 DK127989), and Edward P. Evans Foundation. S.D. is a Scholar of the Blood Cancer United Society (1391-24). This research is also supported by the Fred Hutchinson Cancer Center (FHCC) Shared Resources through NCI Cancer Center Support grants (P30 CA015704 and S10 OD028685).

## AUTHOR CONTRIBUTIONS

E.A.B. designed the study, performed research, analyzed data, and wrote the paper. T-Y.H, A.S., L.B.G., E.A.A-G., E.J.N. performed research and analyzed data. R.V. and S.S., P.B.F., R.S.W. and R.L. provided critical feedback to the study. H.J.D. and D.L.S. provided patient samples. S.D. provided vital reagents, performed research and analyzed data. S.C.L. designed the study, performed research, analyzed data, secured research funding, supervised research, and wrote the paper. All authors read, reviewed and consented to submission of this manuscript.

## Notes

### Competing Interest Statement

The authors have declared no competing interest.

### Summary of Updates

This is an updated manuscript from the initial version. The following changes were made: 1. Title 2. Abstract 3. Manuscript text 4. Minor updates to Figures 2, 3 and 4 4. Order of the supplemental tables

