## Supplemental Figures 1-4 for "CDK8 is a critical effector of cell fate dysregulation in *SF3B1*-mutant MDS"

#### Supplemental Figure Legends

**Supplemental Figure S1. (A)** IGV plots of the skipped exon (SE) between exon 9 and exon 10 in TAF2, Y-axis, absolute read count. **(B)** Normalized read counts (DESeq2) of TAF2 mRNA in *SF3B1* mutant MDS patient samples and cell lines, dashed line marks the mean counts of the *SF3B1* WT patient samples. **(C)** IGV plots of the alternative 3′ splice site (A3SS) between exon 4 and exon 5 in TAF6L, Y-axis, absolute read count. **(D)** Normalized read counts (DESeq2) of TAF6L mRNA in *SF3B1* mutant MDS patient samples and cell lines, dashed line marks the mean counts of the *SF3B1* WT patient samples. **(E)** IGV plots of the A3SS between in exon 17 of MED13L, Y-axis, absolute read count. **(F)** Normalized read counts (DESeq2) of MED13L mRNA in *SF3B1* mutant MDS patient samples and cell lines, dashed line marks the mean counts of the *SF3B1* WT patient samples. **(G)** IGV plots of the A3SS between in exon 5 of MED6, Y-axis, absolute read count. **(H)** Normalized read counts (DESeq2) of MED6 mRNA in *SF3B1* mutant MDS patient samples and cell lines, dashed line marks the mean counts of the *SF3B1* WT patient samples. **(I)** IGV plots of the A3SS in CDK8 exon 8 in AML patient and cell lines with different splicing mutations (*SF3B1* K700E, *SRSF2* P95H, *U2AF1* S34F), Y-axis, absolute read count.

**Supplemental Figure S2. (A)** Diagram of sgRNA's targeting CDK8. **(B)** Analysis (CRISPResso) of editing efficiency surrounding CDK8 guide RNA 2 (sgRNA2). Top – percent sequence difference (green line), Bottom - -log10 of the p value for the edits at each position (blue line). Dashed black line – Bonferroni cutoff correction. Percent sequence difference and p value calculated compared to the reference sequence (ENSG00000132964). Red dashed line – predicted cleavage position. **(C)** Sequence alignments from amplicon sequencing surrounding CDK8 sgRNA2 and percentage of each alignment of total reads. Dashed black line – predicted cut site. **(D)** Representative flow plots from $CD34^+$ UCB cells 24 hours post nucleofection with control (sgAAVS1, brown), or CDK8 (sgCDK8, blue) guides or untreated cells (grey). **(E)** Western blot of $CD34^+$ UCB cells 5 days post nucleofection.

**Supplemental Figure S3. (A)** Experimental overview of xenotransplantation assays. **(B)** Mature human ($mTer119^-hCD45^+$) hematopoietic lineage composition (B cells – $CD19^+$, Myeloid – $CD33^+$, T cells – $CD3^+$) in the peripheral blood of mice over 16 weeks. **(C)** Percent peripheral blood mature human hematopoietic cells at 20 weeks post transplantation. **(D)** Percent mature human hematopoietic cells at 20 weeks in the spleen post transplantation. **(E)** Percent mature human hematopoietic cells at 20 weeks in the bone marrow post transplantation. **(F)** Percent human chimerism (human $CD45^+$) in peripheral blood (red), spleen (green), and bone marrow (blue). **(G)** absolute viable cell number of lineage-negative ($Lin^-$) $CD34^+CD38^+$ cells in the bone marrow at 20 weeks post-transplant. Graphs are plotted as mean ± SD, n= 10 mice per group. **(H)** Representative flow plots and gating strategy for progenitor populations in the bone marrow at 20 weeks ($AAVS^{KO}$ – brown, $CDK8^{KO}$ – blue).

**Supplemental Figure S4. (A)** CDK8 qPCR from $CD34^+$ UCB cells treated with viral particles containing a non-targeting control (shNTC) or CDK8 (shCDK8) shRNAs. Data are plotted as the ratio with 18s RNA, normalized to shNTC. **(B)** Western blot from $CD34^+$ UCB cells treated with shNTC or shCDK8. **(C)** Gating strategy and representative flow plots of UCB $CD34^+$ cells treated with shRNAs. Red box – sorted population. **(D)** Reactome analysis of $CDK8^{KO}$ $CD34^+$ UCB cells, showing significant pathways (Benjamini Hochberg corrected $p< 0.05$). X-axis, normalized enrichment score (NES).

**Supplemental Figure S5. (A)** Sanger sequencing traces from WT (top) and *SF3B1*-mutant (bottom) iPSC-HPCs. Yellow bar highlights the C>A mutation encoding the G742D mutation. **(B)** Normalized read counts (DESeq2) of CDK8 mRNA from published RNA-seq data of *SF3B1*-mutant iPSC-HPCs in erythroid differentiation conditions (n=1/condition, 3 conditions total). Erythroid stage is indicated by point shape (circle = early, triangle = late, square = undifferentiated). Representative flow plots and (**C**) mean ± SD (**D**) of $CD71^+CD235a^-$ and $CD71^+CD235a^+$ iPSC-HPCs on day 8 of erythroid differentiation. (**E**) Representative flow plots of *SF3B1*-mutant iPSC-HPCs in erythro-myeloid differentiation over time. (**F**) Mean ± SD of $CD71^+CD235a^+$ *SF3B1*-mutant iPSC-HPCs transduced with CDK8 cDNA or empty vector (EV) control in erythro/myeloid differentiation over time (n=2). **, $p < 0.01$, ***, $p< 0.001$, Student's t-test.

#### Supplemental Figure S1

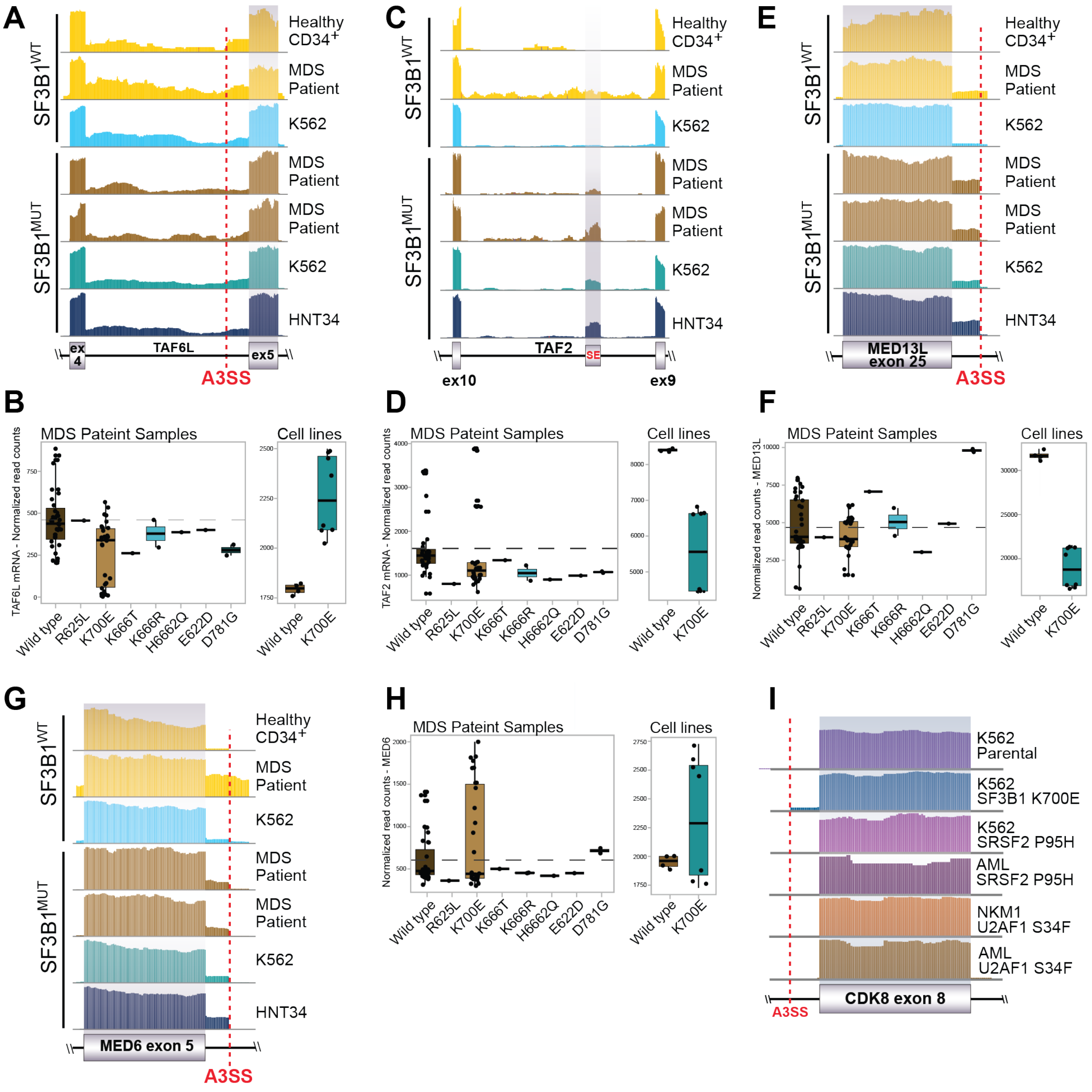

#### Supplemental Figure S2

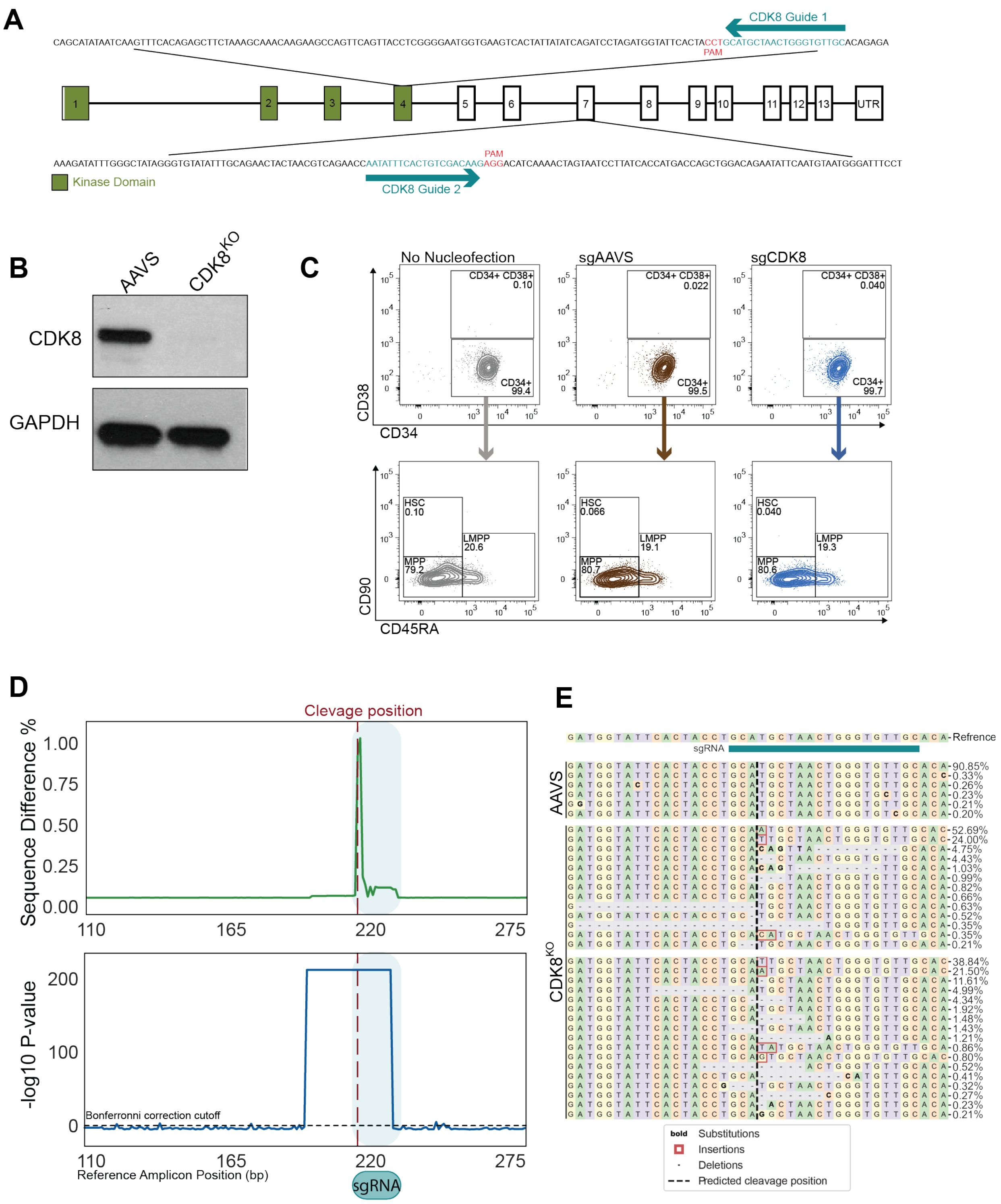

### Supplemental Figure S3

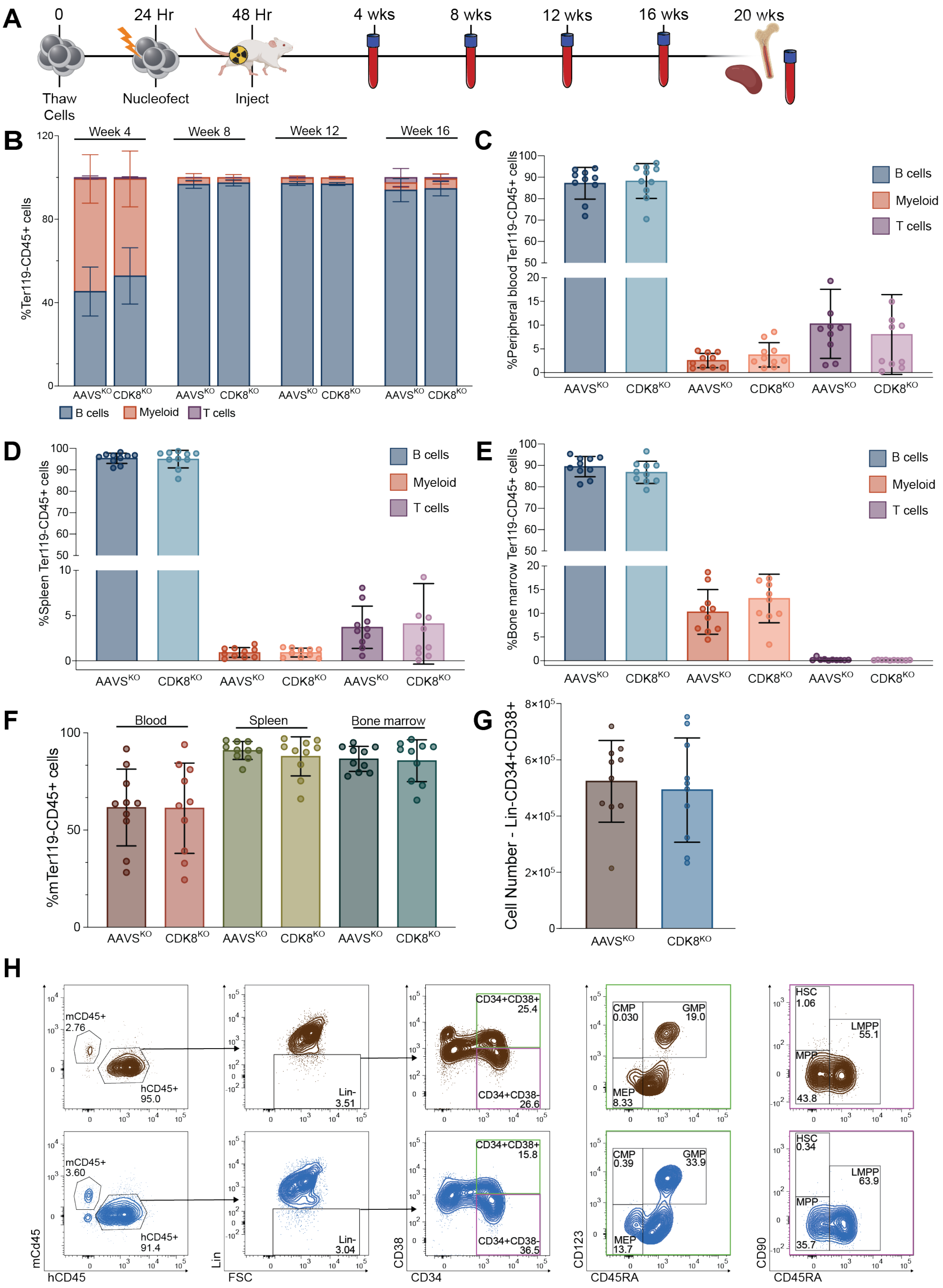

#### Supplemental Figure S4

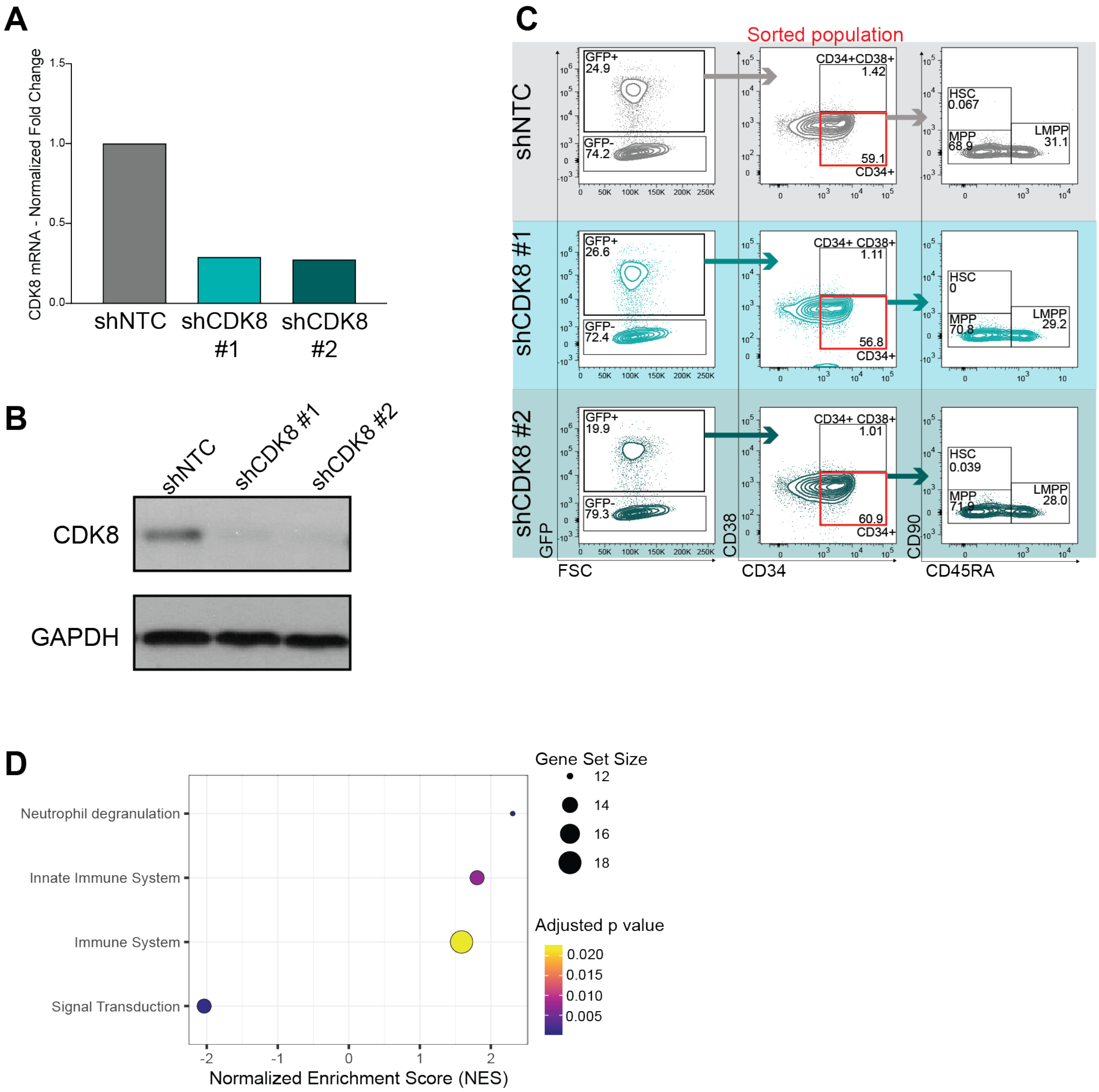

**Supplemental Figure S5**

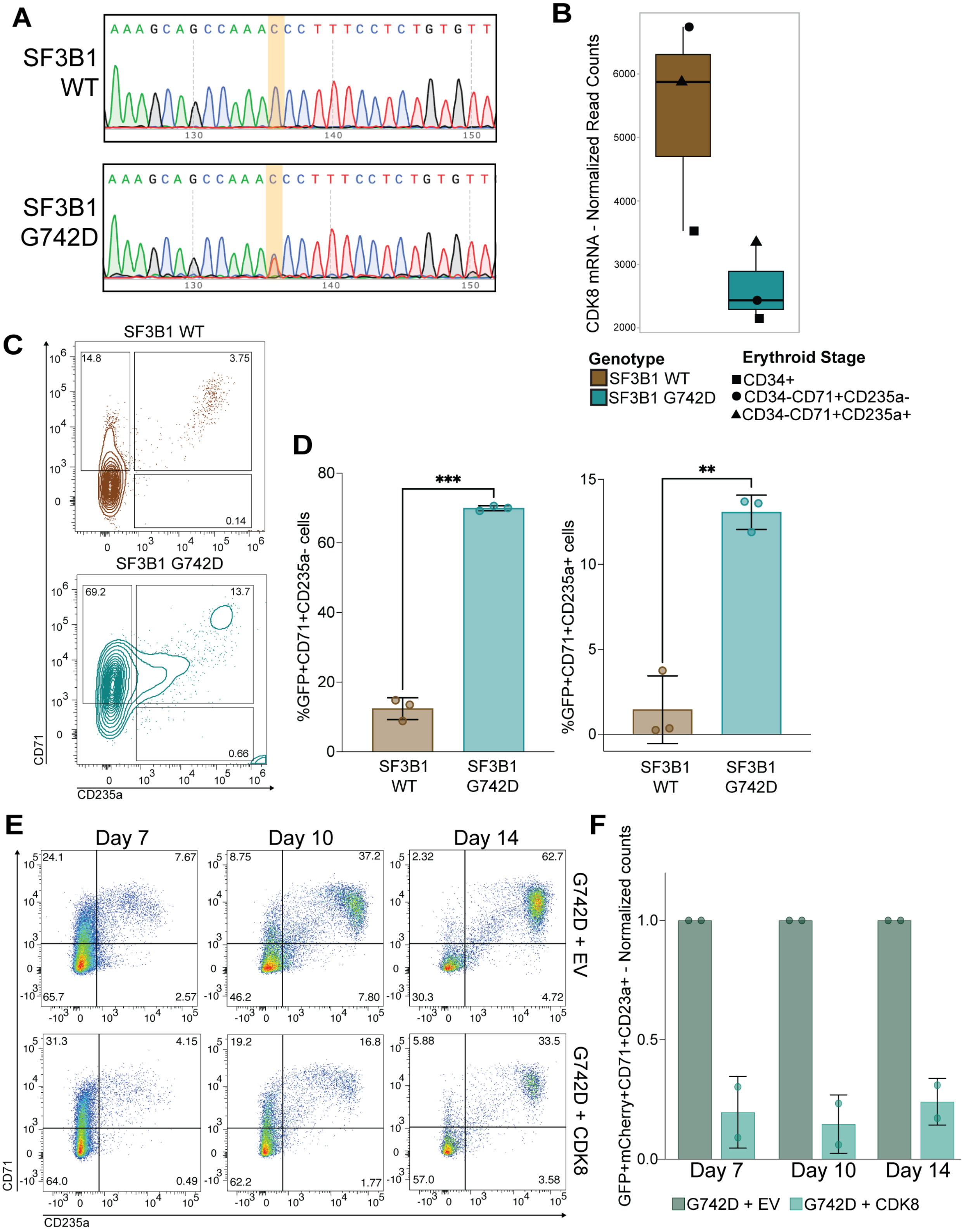
